# Multi-Factor Coral Disease Risk Forecasting for Early Warning and Management

**DOI:** 10.1101/2023.10.23.563632

**Authors:** Jamie M Caldwell, Gang Liu, Erick Geiger, Scott F Heron, C Mark Eakin, Jacqueline De La Cour, Austin Greene, Laurie Raymundo, Jen Dryden, Audrey Schlaff, Jessica S Stella, Tye L Kindinger, Courtney S Couch, Douglas Fenner, Whitney Hoot, Derek Manzello, Megan J Donahue

## Abstract

Ecological forecasts are becoming increasingly valuable tools for conservation and management. However, there are few examples of near real-time forecasting systems that account for the wide range of ecological complexities. We developed a new coral disease ecological forecasting system that explores a suite of ecological relationships and their uncertainty and investigates how forecast skill changes with shorter lead times. The Multi-Factor Coral Disease Risk product introduced here uses a combination of ecological and marine environmental conditions to predict risk of white syndromes and growth anomalies across reefs in the central and western Pacific and along the east coast of Australia and is available through the U.S. National Oceanic and Atmospheric Administration Coral Reef Watch program. This product produces weekly forecasts for a moving window of six months at ∼5 km resolution based on quantile regression forests. The forecasts show superior skill at predicting disease risk on withheld survey data from 2012-2020 compared with predecessor forecast systems, with the biggest improvements shown for predicting disease risk at mid-to high-disease levels. Most of the prediction uncertainty arises from model uncertainty and therefore prediction accuracy and precision do not improve substantially with shorter lead times. This result arises because many predictor variables cannot be accurately forecasted, which is a common challenge across ecosystems. Weekly forecasts and scenarios can be explored through an online decision support tool and data explorer, co-developed with end-user groups to improve use and understanding of ecological forecasts. The models provide near real-time disease risk assessments and allow users to refine predictions and assess intervention scenarios. This work advances the field of ecological forecasting with real world complexities, and in doing so, better supports near term decision making for coral reef ecosystem managers and stakeholders. Secondarily, we identify clear needs and provide recommendations to further enhance our ability to forecast coral disease risk.

## Introduction

Forecasting coral disease outbreaks is critical for the timely management of reef ecosystems, but developing such early warning systems is challenging when disease dynamics are not well understood, and data are sparse and irregularly updated. Coral reefs and their associated threats are a prime example of a complex system that presents challenges for ecological forecasting. Coral reefs are dynamic, heterogeneous environments, characterized by their diversity of species and species interactions, and physical and chemical factors that can affect their health. Diseases are a major threat to coral reefs, causing up to 95% mortality in dominant coral species during outbreak events such as the white band disease epidemic in the 1980s and 1990s and the Stony Coral Tissue Loss Disease outbreak in the 2010s and 2020s in Florida and the Caribbean (Aronson and Precht 2001; Walton et al. 2018; Rosales et al. 2020). Thus, innovative approaches to support effective management strategies for coral disease transmission, prevention, and mitigation are urgently needed. Modern forecasting can now combine mechanistic understanding, statistical and machine learning models, and the quantification of uncertainty to make more accurate predictions. However, developing accurate and reliable forecasts requires overcoming several key challenges, including the sparse availability of high-quality data and limited understanding of the underlying complexity of biological and environmental drivers of coral disease outcomes.

Developing early warning systems for coral diseases is a relatively recent endeavor that aims to help managers and decision-makers take preventative actions and mitigative measures. Given the widespread consensus linking coral disease to thermal condition (Burke et al. 2023), early disease forecasts focused on using temperature to predict suitable conditions for disease. The U.S. National Oceanic and Atmospheric Administration Coral Reef Watch (NOAA CRW) program developed the first coral disease forecast in 2010 for white syndromes on the Great Barrier Reef (GBR) in Australia (hereafter V1), which uses a decision tree framework based on a series of anomalous thermal metrics (Heron et al., 2010). This system leveraged NOAA CRW’s established satellite sea surface temperature (SST) monitoring data (∼50 km resolution, twice-weekly) refining previous information about the relationship between thermal condition and disease (Bruno et al., 2007; Selig et al., 2006). Subsequently, NOAA CRW adapted the thermal condition metrics from the GBR to produce a complementary, experimental predictive tool for coral disease in the Hawaiian archipelago (incorporated into V1). V1 was further developed to incorporate finer resolution (∼5 km, daily) SST data (hereafter V2). A retrospective analysis of V2 demonstrated that machine learning algorithms that used the product metrics (i.e., summer Hot Snaps) combined with additional biotic data could robustly reproduce disease prevalence patterns for two coral diseases across three host species (Caldwell, Heron, et al., 2016). The ability to nowcast and forecast some of these reef stressors has led to new and innovative conservation practices and provided clarity to managers seeking to set priorities. While we know of no examples yet where management officials have taken action in response to disease forecasts, we have seen responses to NOAA CRW bleaching forecasts (Raymundo et al., 2022) and we expect complementary actions (Beeden et al., 2012; Neely et al., 2021) will be taken in response to disease forecasts as managers become more familiar with the system. Managers could mitigate disease risk and impacts with a variety of local scale actions such as implementing fishing and fishing gear restrictions, reducing land-based pollution runoff, or reducing the abundance of known disease vectors (eg., corallivorous gastropods) or predators (e.g., Acanthaster planci) from vulnerable reefs. Other experimental approaches could be effective, including probiotics, phage therapy, or temporarily relocating at-risk colonies to aquaria.

Moving beyond thermal conditions, the next generation of coral disease early warning systems needs to better incorporate an expanded suite of conditions known or hypothesized to affect disease dynamics. Previous research has statistically linked a range of conditions with impaired coral health, including colony size and density, thermal condition, water quality, human population density and land use, and fish densities and predation (Aeby, Williams, Franklin, Haapkyla, et al., 2011; Aeby, Williams, Franklin, Kenyon, et al., 2011; Bruno et al., 2003; Caldwell et al., 2020; Carlson et al., 2019; Greene et al., 2020; Haapkylä et al., 2011; Pollock et al., 2014; Redding et al., 2013; Renzi et al., 2022). However, the mechanistic underpinnings of these multiple contributing factors are often poorly understood due to their complex and non-linear behavior, which can vary by host species and disease type (Clemens & Brandt, 2015; Shore & Caldwell, 2019; Vega Thurber et al., 2014). An additional challenge for any early warning system is whether the predictor variables themselves can be forecasted (Clark et al., 2001; Oliver & Roy, 2015), and this is especially true for the diverse drivers of coral disease. These challenges must be addressed to incorporate a wider range of putative disease drivers into forecasting models.

Over the last decade, ecological forecasting and monitoring tools have advanced considerably, making it possible to integrate multiple data streams and more robustly consider various scenarios and sources of uncertainty (Clark et al., 2001; Dietze et al., 2018). Machine learning algorithms are particularly useful in this context, as they can identify complex non-linear relationships between variables and make predictions in data-poor environments (Jordan & Mitchell, 2015). These advancements have enhanced the capability of identifying relationships that should be tested further, allowing incremental improvements in forecasting efforts (Dietze et al., 2018). Exploring likely scenarios within a forecasting framework can help create more robust approaches for managing ecosystems. Plausible scenarios with alternate conditions are developed using a combination of scientific information and models, stakeholder input, and expert opinion. Scenarios can then be used to explore the potential impacts of different management strategies, account for additional spatial variation in predictor variables, and/or refine predictions to a specific set of conditions. By incorporating scenarios into ecological forecasts and management plans, managers and decision-makers can better understand the potential outcomes of different decisions and identify strategies that are more likely to be effective under a range of possible futures (Clark et al., 2001). These models also need to be incorporated into easy-to-use tools for managers to test and compare different management actions.

In this paper, we present the next-generation NOAA CRW coral disease forecasting product (i.e., V3) that addresses some of these challenges and applies new, innovative approaches to ecological forecasting. By integrating data from multiple sources and using machine learning algorithms to identify patterns and make predictions, the system provides early warnings of coral disease risk and could help managers and decision-makers take proactive measures to protect reefs across much of the Pacific Ocean. This new Multi-Factor Coral Disease Risk product expands the previous product in several ways through: 1) a broader geographic scope; 2) consideration of two distinct groups of diseases; 3) inclusion of a suite of disease drivers; 4) generation of weekly forecasts with up to three-month lead time; 5) provision of measures of uncertainty; 6) consideration of multiple scenarios; and 7) capacity for users to visualize forecasts and modify scenarios through an interactive online dashboard used to explore management strategies. The results of this study have broader implications for making predictions in other complex, data-poor systems and highlight the need for continued research and innovation in the field of ecological forecasting.

## Methods

The Multi-Factor Coral Disease Risk product (i.e., V3) is an experimental regional product, currently providing ecological forecasts for multiple locations in the Pacific Ocean. Areas include American Samoa, Guam and the Commonwealth of the Northern Mariana Islands (CNMI), Australia’s Great Barrier Reef (GBR), Hawaii, and the U.S. Pacific Remote Islands Marine National Monument (PRIMNM, also called the Pacific Remote Island Area, PRIA) encompassing seven islands and atolls: Baker, Howland, and Jarvis Island; Johnston, Wake, and Palmyra Atolls; and Kingman Reef. In this product, we assess disease risk based on satellite remotely-sensed, modeled, and *in situ* data to provide nowcasts and near-term forecasts based on current conditions, recent conditions, and subseasonal-to-seasonal forecasts from NOAA operational climate models. We defined disease risk separately as a density (number of diseased colonies/75m2 ranging from 0 to infinity) in Australia and as a prevalence (percent of colonies affected ranging from 0 to 100%) in the U.S. Pacific (more details below), which maps to different NOAA CRW warning levels ranging from Low Risk to Alert Level 2 for visualization purposes in the decision support tool. We determined the thresholds separating warning levels based on historical disease observations and expert elicitation; the thresholds vary by disease type and region (Appendix S1: Table S1). We optimized the modeling system using a Pacific-wide dataset of over 42,000 coral disease surveys (more detail below).

## Data

We identified a suite of potential predictor variables to forecast coral disease risk based on prior observational, experimental, and modeling efforts for two disease types: white syndromes and growth anomalies (Table 1, Appendix S1: Table S2).. White syndromes refer to a suite of tissue loss diseases that cause coral mortality and range from acute to chronic based on the speed at which the infection progresses (Bourne et al., 2015). Growth anomalies are chronic diseases that persist at low levels year round and manifest as changes in skeletal morphology, usually through abnormal increases in skeleton secretion and disorganization of corallites, affecting colony growth and fecundity (Palmer & Baird, 2018). The etiological agents of both groups of diseases are unknown. Across a variety of host species, disease types, and regions, some factors such as coral cover, coral colony size, and specific ranges of temperature have been consistently associated with certain coral diseases, although the functional relationships may differ slightly (Bruno et al., 2007; Caldwell et al., 2020; Greene et al., 2020; Heron et al., 2010; Ruiz-Moreno et al., 2012). Thus, we considered appropriate derivations of these variables for all diseases and regions, based on data availability. For instance, we considered accumulation of anomalous temperatures for white syndromes because of statistical associations across multiple large scale studies (Bruno et al., 2007; Burke et al., 2023; Heron et al., 2010; Howells et al., 2020; Maynard et al., 2011), but focused on seasonal mean temperature for growth anomalies because it has been experimentally associated with lesion development and growth (Stimson, 2011). Additionally, there were several potential predictor variables that were unique to each disease type, because the ecologies of white syndromes and growth anomalies differ substantially. White syndromes often exhibit strong seasonality due to ocean conditions, notably winter and summertime thermal stress and changes in water quality (Haapkylä et al., 2011; Heron et al., 2010; Maynard et al., 2011; Ruiz-Moreno et al., 2012). White syndromes have also been associated with fish densities, but the effects are not consistent across studies and the underlying hypothesis for this effect varies by fish functional group (Aeby, Williams, Franklin, Kenyon, et al., 2011; Caldwell et al., 2020; Clemens & Brandt, 2015; Greene et al., 2020; Renzi et al., 2022; Williams et al., 2010). Thus, we included available metrics of fish density for multiple fish types and water quality (turbidity) in the white syndromes models. For growth anomalies, previous studies indicate an association with low fish abundance, limited water motion, and poor water quality via nutrient enrichment, coastal development, and proximity to dense human populations (Aeby, Williams, Franklin, Haapkyla, et al., 2011; Caldwell et al., 2020; Yoshioka et al., 2016). Therefore, we included metrics of fish populations, turbidity, and coastal development in the growth anomalies models.

**Table 1:** Variable inclusion and importance differs for each disease-region model. The variables tested and selected, as well as their importance, differ for each region (Great Barrier Reef or U.S. Pacific) and disease type (white syndrome or growth anomalies). A cell with a value indicates that the variable was selected for the model and the value represents the percent increase in Mean Squared Error (MSE) of out-of-bag cross-validation predictions across permutations in that predictor variable, with higher values indicating more important predictor variables. (Note that MSE is sensitive to units even though the percent increase in MSE is unitless; thus values for the GBR models that predict disease density will typically be much larger than values for the U.S. Pacific models that predict disease prevalence). x indicates a variable was tested but not selected; a blank cell indicates that the variable was not tested for that model because it is not a hypothesized predictor variable whereas ∅ indicates that a variable was not tested because data were not available. Metrics that measure Kd(490) are a proxy for turbidity.

| Predictor variable | White syndromes |  | Growth anomalies |  |
| --- | --- | --- | --- | --- |
|  | GBR | U.S. Pac | GBR | U.S. Pac |
| *Time-invariant predictors* |  |  |  |  |
| Coral cover | 51 | 0.8 | 487 | x |
| Median colony size | ∅ | 2.3 | ∅ | 5.3 |
| Colony size variability |  |  | ∅ | x |
| Herbivorous fish density | 68 | 1.2 | 641 | 2.3 |
| Parrotfish density | ∅ | 0.6 |  |  |
| Butterflyfish density | ∅ | x |  |  |
| Long term Kd(490) median | x | 2.2 | x | 1.9 |
| Long term Kd(490) variability | x | 1.9 | 399 | x |
| Coastal development |  |  | x | 3.1 |
| *Seasonally-changing predictors* |  |  |  |  |
| Three-week Kd(490) median | x | 1.3 | 350 | 1.8 |
| Three-week Kd(490) variability | 59 | 1.7 | 345 | x |
| Month | 37 | 1.1 | 310 | 2.5 |
| *Regularly-changing predictors* |  |  |  |  |
| 90-day SST mean |  |  | 413 | 5.6 |
| Hot Snap | 53 | x |  |  |
| Winter Condition | 62 | 0.8 |  |  |

From a forecasting perspective, the predictor variables, or environmental conditions, that we considered in this study can be roughly divided into three types based on their variabilities through time: 1) time-invariant; 2) seasonally-changing; and 3) regularly-changing. We consider time-invariant conditions as any predictor variable that does not change regularly through time, or information about such change is unavailable or sparsely updated. We consider seasonally-changing conditions as predictor variables that depend on time of year but are not date-specific. Most of these variables have been developed in a way that represents repeated seasonal patterns developed from multi-year datasets (i.e., climatologies). Finally, we consider regularly-changing conditions as predictor variables that change, and can be measured and evaluated, over some regular time interval. We used point estimate predictor variable data for model development based on the time and location of coral disease surveys whereas we use gridded predictor variable data for forecasts.

### Time-invariant data

We collated time-invariant data from *in situ* surveys and remotely-sensed data. For each predictor variable described below, which we used in at least one of the four region-by-disease models, we provide further detail, including data sources and spatial resolution, in Appendix 1: Table S2. To characterize benthic cover, we used metrics of coral cover (0-100%), coral colony size, and population level colony size variability (based on coefficient of variation). For these metrics, we developed the models using data collected concurrently with coral disease surveys. We aggregated these metrics by host family for both U.S. Pacific models and by morphology for the GBR white syndromes models to be consistent with data collection methodology (more details below). In the forecasts, we used a combination of survey data and gridded data from long-term monitoring programs. For coral cover in the GBR and coral colony size in the U.S. Pacific, we calculated ∼5 km pixel-specific mean values across the reef grid from long-term survey data (multiple sources listed in Appendix 1: Table S3) while for coral cover in the U.S. Pacific, we used sector level benthic cover data from the NOAA National Coral Reef Monitoring Program (NCRMP). As fish surveys were rarely conducted in coordination with benthic surveys, we used fish density layers from long-term monitoring programs for both model development and forecasting. We used sector-level fish data from NOAA NCRMP and ∼2 km gridded fish count data based on manta tow surveys from the Australian Institute of Marine Science (AIMS) Long Term Monitoring Program (LTMP) (Sweatman et al., 2008). These long-term datasets represent the most comprehensive current estimates of coral cover and size available for the reef grid, but inherently will not contain information about recent or future changes in these variables. Thus, periodic updates to the reef grid as data becomes available would be beneficial. As a proxy for coastal development in both model development and forecasting, we used NASA’s Black Marble product, which is a time-aggregated map of artificial light intensity (high gain) (range = 0-255 where 0 = black and 255 = white) at 3 km resolution from the Visible Infrared Imaging Radiometer Suite (VIIRS) instrument aboard the Suomi-National Polar-orbiting Partnership (NPP) satellite (Román et al., 2018). To characterize chronic water quality conditions in both model development and forecasting, we aggregated the diffuse attenuation coefficient at 490 nm, Kd(490), as a proxy for turbidity from VIIRS data (Kirk, 1994). We calculated long-term Kd(490) median and variability for each reef pixel by overlaying aggregated data from 2012-2020 (i.e., all data available at the time of study) within a 5-pixel buffer (750 m becomes ∼8.25 km resolution) following methods from Geiger et al., 2021 to increase data availability, as nearshore ocean color data are notoriously patchy. These metrics are indicative of spatial differences in water quality across reefs, providing information on locations that have chronically good or poor water quality and those that are exposed to a large range of water quality conditions throughout the year versus those with more consistent conditions.

### Seasonally-changing data

We use month of year and two turbidity metrics (mean and variability) to capture seasonally-changing conditions that are relevant to disease risk. To characterize typical seasonal water quality patterns, we calculated mean and variability of VIIRS-derived Kd(490) for a three-week moving window (resulting in new values each week) across a 9-year time span (2012-2020) using the same 5-pixel buffer described above. These metrics repeat annually and are indicative of how water quality changes throughout the year at a given location. We used the mean (i.e., climatology) and associated variability in Kd(490) to represent seasonal changes because, to date, these values are too highly variable and too infrequently available in the coastal zone to use actual, or even three-week composite, values. Additional details on the derivation of these metrics and their accuracy can be found in Geiger et al., 2021. We include month in the model as a proxy for all other seasonally changing conditions.

### Regularly-changing data

We include three temperature-based metrics in the disease models that update at regular intervals: 90-day SST mean, Hot Snap, and Winter Condition. In contrast to the seasonally-changing data, the regularly changing data yield different values each year for the same time period (e.g., the first week of January) based on observed and/or forecasted conditions. The daily previous 90-day mean SST is the average daily SST values for the 90 days preceding the current date. The Hot Snap and Winter Condition metrics were developed for NOAA CRW’s Coral Disease Outbreak Risk Product V1 and V2 and continue to be used in V3 to represent thermal conditions on time scales relevant to coral disease. The Hot Snap metric accumulates hot temperature anomalies through time, relative to the locally/pixel-specific long-term expected summer season temperature (summer season climatology) (Heron et al., 2010), providing an indication of exposure to thermal stress. The Winter Condition metric accumulates both hot and cold temperature anomalies during the winter season relative to locally-specific, long-term average temperature (Heron et al., 2010), representing cold season variability. Mild winters have been linked to white syndromes (Caldwell, Heron, et al., 2016; Heron et al., 2010), potentially because such conditions allow pathogens to persist and grow throughout the winter season.

The data underlying these three temperature metrics differ for satellite observed temperatures, which we used for model development and near real-time nowcasts, and forecasted temperature, which we use for disease forecasts. For observed SST, we use CoralTemp v3.1 (Skirving et al., 2020), which provides daily data at a global resolution of ∼5 km (0.05°). For forecasted SST, we use output from the NOAA National Centers for Environmental Prediction’s operational Climate Forecast System Version 2 (CFSv2) (Saha et al., 2014). Each day, the CFSv2 generates an ensemble of four SST forecasts out to 9 months at ∼50 km (0.5°) resolution. We use this output to form 28 ensemble member predictions each week (4 daily start times x 7 days) for each predicted temperature metric and to predict the metric for each future week up to three months following the prediction date. The data are downscaled from ∼50 km to ∼5 km using a nearest neighbor algorithm and then bias-corrected to match the 5 km satellite SST measurements during the overlap between satellite observations and CFSv2 over the weekly time period when the 28 ensemble members are collected. Because predicted values demonstrate decreased variability with longer lead-times, predicted SST anomaly values for the metrics are correspondingly adjusted to match the variability of the SST data.

### Coral Disease Survey Data

We assembled a Pacific-wide coral health monitoring dataset, which we used to develop region- and disease-specific predictive models of disease risk. In total, we assembled over 42,000 coral health monitoring surveys between 2012 and 2020. Data came from the NOAA NCRMP, University of Guam, Hawaii Coral Disease Database (Caldwell, Burns, et al., 2016), and the Great Barrier Reef Marine Park Authority (GBRMPA; referred to as Reef Authority in other contexts) Reef Health and Impact Surveys (Beeden et al., 2014). The different survey protocols used to collect these data have been described in detail previously (Beeden et al., 2014; Caldwell, Burns, et al., 2016; Winston et al., 2020). For the research described in this paper, there are notable methodological differences between surveys conducted in Australia and the U.S. Pacific; therefore, we modeled disease risk in these two regions separately. Specifically, surveys in Australia indicated morphology-specific disease density (i.e., number of diseased colonies in a given area) while the U.S. Pacific surveys provided information to quantify family-specific disease prevalence (i.e., percent of coral colonies affected by disease); therefore the risk prediction is for disease density for Australia and disease prevalence in the U.S. Pacific. While the U.S. Pacific models are technically generated at the family level (Acroporidae for white syndromes and Poritidae for growth anomalies), in practice, the data predominantly describe genus- or species-specific patterns with various genera/species represented in different regions (Appendix 1: Table S4). We used these data to develop predictive models of disease risk (i.e., disease density or prevalence) rather than outbreak risk, which we believe is more appropriate as the data arise from regular monitoring surveys rather than outbreak response surveys (outbreaks defined in Raymundo et al., 2008). If multiple surveys were conducted in close proximity in time (i.e., in the same month) and space (the same survey area), we randomly selected one of those surveys to keep in the dataset to avoid artificially over-representing certain conditions.

### Balancing data with SMOTE

To create disease models that produce reliable predictions of all levels of disease risk, particularly of high disease levels, we used a synthetic sampling technique to balance the data used in model development. The observational surveys available to create the disease models were highly unbalanced, with the majority of surveys reporting zero or low levels of disease (Appendix 1: Table S5); using unbalanced data would optimize disease-free predictions.

Therefore, we balanced the dataset before model creation using the Synthetic Minority Over-sampling Technique (SMOTE; Chawla et al., 2002; Fig. 1A). We used the observed disease surveys with their associated predictor variables to create additional synthetic disease surveys (using a k-nearest neighbor algorithm) to produce a balanced dataset (e.g., approximately equal number of disease and disease-free surveys with all predictor variables). We created multiple SMOTE datasets for each disease and region based on different disease level thresholds because it is unknown whether the same environmental conditions that precede observed disease are responsible for low and high levels of disease risk. In other words, we oversampled surveys with any disease and oversampled surveys with greater than some specified level of disease allowing the model selection process to determine the best threshold to use. We tested several disease level thresholds: 1, 5, and 10 colonies/75m^2^ for white syndromes in the GBR and 1, 5, and 10% prevalence for white syndromes in the U.S. Pacific; 1, 5, 10, and 15 colonies/75m^2^ for growth anomalies in the GBR and 1, 5, 10, 15, and 20% prevalence for growth anomalies in the U.S. Pacific. We chose these thresholds based on a combination of natural breaks in the data and expert opinion. The threshold units align with the survey data collected (i.e., density or prevalence) and therefore differ between the GBR and U.S. Pacific. For each SMOTE dataset of disease surveys, we split the data into training and test data using a 75/25 split. We used the training data for model creation and then the withheld test data for model selection and assessment (described below).

**Fig 1.**
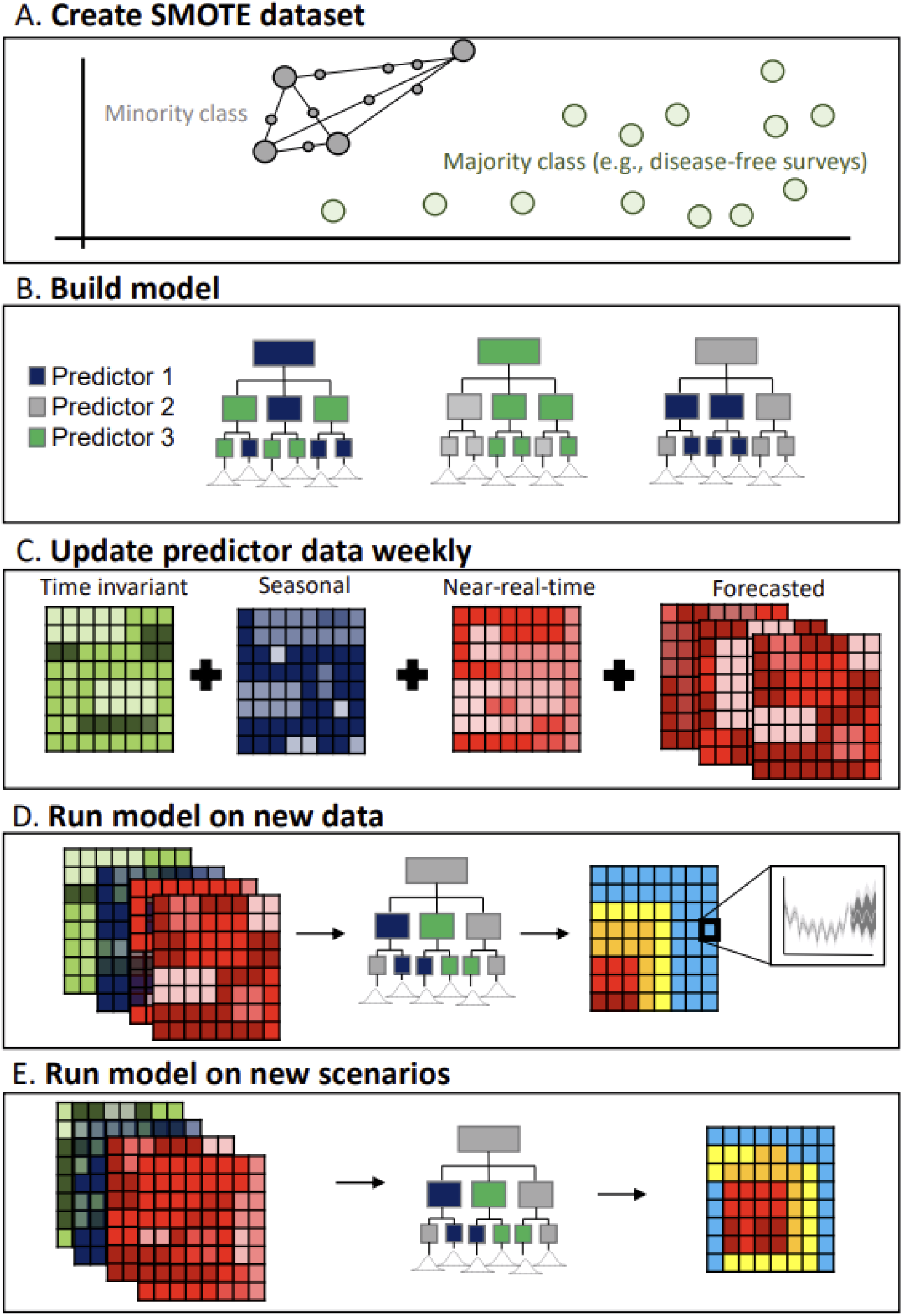
Methodological overview for model development and weekly update for each disease-by-region model. A) Graphical illustration of Synthetic Minority Over-sampling Technique (SMOTE) where the minority class (i.e., surveys with disease; large gray circles) are used to create synthetic surveys of predictor and response data (i.e., small gray circles) based on k-nearest neighbors (i.e., black lines connecting surveys in n-dimensional parameter space), resulting in approximately equal numbers of surveys with (gray) and without (green) disease present. In this study, we tested different thresholds for inclusion in the minority class. B) We built the model using quantile regression forests, an algorithm that creates many decision trees based on a subsample of predictor variables (example shows each tree using 2 of 3 possible predictor variables), and produces a distribution of target values rather than a mean value. We selected the most parsimonious model across the different SMOTE datasets and quantile regression forests (i.e., with different combinations of predictor variables) based on a withheld portion of the data, using the models with the fewest number of predictor variables with superior predictive skill. The selected models are used in the weekly update for C-E. C) Each week, we update predictor variables for the reef grid. Time-invariant predictor variables are held constant, seasonal predictor variables update each week or month, near-real-time data reflect recent satellite observations, and forecasted data come from 28-member ensemble CFSv2 SST forecasts. D) Using the updated predictor data, we re-run the model to produce a new near real-time prediction and 12 weeks of forecasted data, which we amend to the prior 11 weeks of historical nowcast predictions for a total of six continuous months of disease risk assessments. E) We also vary the predictor data across a gradient of values to produce scenarios, to explore how disease risk changes with different input variable values.

### Quantile regression forests

We created predictive models of disease risk using quantile regression forests (Meinshausen, 2006; Fig. 1B). Quantile regression forests use a decision tree framework, allow for non-linear relationships between response and predictor variables, and have shown high predictive skill across a range of systems. Briefly, quantile regression forests are created by developing an ensemble of quantile decision trees (i.e., random forests), with each tree created from a bootstrapped resample of the dataset. Quantile decision trees differ from standard decision trees in that they predict the distribution of target values rather than the mean target value from the training data. This approach uses an ensemble of uncorrelated decision trees, which tend to outperform any individual tree, and each tree uses a random subset of predictors to increase variation among trees. The result of this process is that the final predictive model is more robust because it is created from many trees that are trained on different subsets of response data and predictor variables.

### Model selection

We selected the most parsimonious model for each disease and region amongst a suite of candidate models based on predictive skill on a withheld portion of the data. For each disease-by-region pair, we ran a model that included all hypothesized predictor variables (Table 1) from a training dataset (75% of surveys) and then used a backward selection approach to iteratively remove predictor variables of least importance. We calculated the relative importance of each predictor variable as the percent increase in Mean Squared Error (MSE) of out-of-bag cross-validation predictions across permutations in that predictor variable, with higher values indicating more important predictor variables. The exception was for the predictor variable Month, which we retained in the model regardless of its relative importance because it captures additional seasonal variation. At each model iteration, we predicted disease risk from a withheld test dataset (25% of the surveys) and assessed predictive skill based on the R^2^ value that arose from linearly regressing those predictions with observations. We followed this approach of backward selection for each SMOTE dataset. The selected (most parsimonious) model was the model with the fewest predictor variables that produced an R^2^ value within 1% of the best model (i.e., model with the highest R^2^ overall).

### Model assessment

To determine how well the models performed at retrospectively predicting disease risk for each disease-by-region pair, we compared retrospective predictions by the models described here with archived nowcasts from previous versions of the models where available (i.e., V2 predictions for the GBR and Hawaiian archipelago) and how forecast skill changes with different lead times. For both assessments, we quantified predictive skill using the withheld test data. To assess predictive skill for white syndromes, we compared retrospective disease predictions from models described in this paper (V3) with models supporting V2 using predictor data available from the corresponding week of observations. The V3 models predict disease density or prevalence whereas the V2 models produce risk levels based on Hot Snap values (units = ℃-weeks, range = 1-15); therefore, we visually compared these results but did not directly compare their skill quantitatively. Since there are no previous models in production for growth anomalies, we assessed the retrospective model skill on the withheld test data alone. Additionally, we were interested in whether and to what extent forecast prediction accuracy and precision change as we get closer to the observation date (i.e., shorter lead-times). To assess this relationship, we predicted disease risk at weekly intervals for each observation date in the withheld data, with lead times ranging from 12 weeks prior (e.g., in advance of a survey) to 0 weeks (i.e., nowcast). We calculated accuracy as the difference between the 75th quantile prediction and the observation, resulting in zero if there was perfect accuracy, negative values if the models predicted lower disease risk than observed, and positive values if the models predicted higher disease risk than observed. We used the 75th quantile prediction (upper range of disease likelihood) as the primary indicator of disease risk throughout this work, which was the metric selected by the product end users to err on the side of potentially overpredicting disease in an effort to further capture rare disease events. To assess predictive precision, we calculated the difference between the 90th and 50th quantile predictions: larger differences indicate less precise estimates and smaller differences indicate more precise estimates.

### Weekly-updating predictions

The overarching objective of this research was to develop a product that provides weekly-updated, near real-time, and subseasonal-to-seasonal disease risk forecasts. The workflow for this process follows. First, we developed a reef location database based on a ∼5 km gridded reef locations dataset currently used by NOAA CRW (Heron et al., 2016) to set the spatial extent of the disease risk forecasts described in this paper. This reef location database encompasses all known shallow-water reefs within the U.S. Pacific Islands and atolls and along the east coast of Australia, the majority of which fall within the GBR Marine Park. To allow users to assess short-term temporal evolution of disease risk at each reef pixel, we provide a moving window of six months of weekly predictions: the first three months with weekly nowcast predictions based on observed environmental conditions (i.e., time-invariant, seasonally-changing, and nowcast predictor variables identified in the model selection process described above) up to the current calendar week, and the second three months with weekly forecast predictions based on a combination of historically observed (time-invariant data and seasonally-changing data) and forecasted environmental conditions. The models and environmental conditions we use vary by disease and region, as described earlier and in the Results section. For each week of predictions, we update the environmental input data (Fig. 1C). The nowcast predictions (Fig. 1D) that we produce for each reef pixel are based on a single set of observed environmental conditions and prediction uncertainty arises solely due to model uncertainty. In contrast, we produce 28 ensemble forecast predictions for each reef pixel (Fig. 1D), using 28 sets of SST-based metrics derived from the 28 different CFSv2 model runs, and thus, uncertainty is composed of both model uncertainty and SST forecast uncertainty. In this product, we chose to present predictions using the 50th, 75th, and 90th quantile predictions for the reasons stated above (though any quantile(s) could be used). We also aggregate the risk predictions for different management areas, which we collated from marine management agencies. We do this by quantifying the 90th quantile values across all ∼5 km reef pixels that fall within the specified management area of the risk predictions (i.e., the 75th quantile modeled risk). The use of the 90th quantile to spatially summarize risk predictions is consistent with other regional summaries produced by CRW (Heron et al., 2016), with this value selected to alert users to regional-level risk whilst preventing potential exaggeration (e.g., by reporting the maximum value across the region).

### Weekly-updated scenarios

To allow users to customize the prediction to localized and current environmental conditions and help determine the most appropriate intervention strategies, we also produce weekly-updated scenario-based disease risk predictions (Fig. 1E). The predictions for various scenarios show how adjusting current environmental conditions would change current disease risk predictions. We calculate the change in disease risk by re-running the models iteratively, varying a single environmental condition by specified amounts and holding all other location-specific, current environmental conditions constant. The resulting scenarios allow users to 1) refine predictions considering local conditions (e.g., a reef of interest) known to the user that may vary from the mean conditions of the entire reef pixel or management zone; and 2) consider how an intervention (e.g., a program to reduce turbidity) would affect disease risk. Following the format we use to present near real-time and seasonal disease risk predictions, we also calculate changes in disease risk for scenarios based on the 75th quantile disease predictions and aggregate the results to management areas in the same way we describe earlier.

## Results

### Performance evaluation

The new Multi-Factor Coral Disease Risk product (V3) described in this study predicts disease risk relatively well and qualitatively demonstrates superior predictive accuracy compared with V2 for both the GBR and Hawaiian archipelago (Fig. 2). All versions have difficulty predicting no or very low disease levels (i.e., below the selected SMOTE thresholds). V3 is the first product to calculate uncertainty and can therefore represent this lack of predictability with large uncertainty values, as shown around many low disease values. The major improvement can be seen at mid- and high-levels of disease (i.e., above the selected SMOTE thresholds, which vary by disease and region). While the previous algorithm predicted some high disease events well, many were predicted to have no disease risk, suggesting that factors other than thermal condition are key for predicting disease events.

**Fig. 2.**
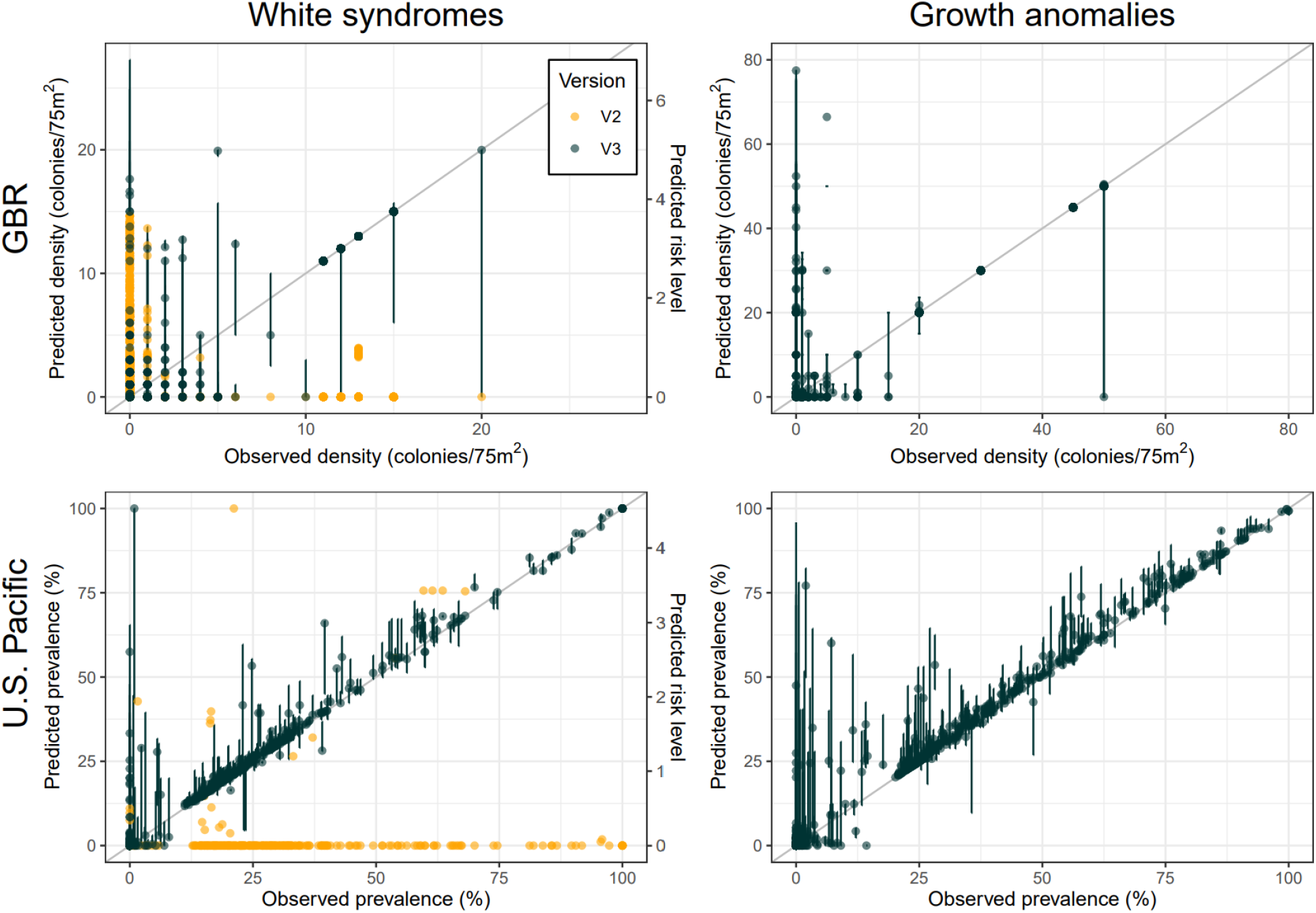
Accuracy of disease nowcast predictions demonstrate improved predictive capabilities for V3 compared with its predecessor. We show comparisons of disease observations (x-axes) with disease predictions (y-axes) for the current model (V3). Points that fall on the gray line indicate a perfect fit between observations and predictions. For white syndromes (left column), we compare disease predictions from V3 with V2 (note that predictions are only available for Hawaii in the U.S. Pacific). For growth anomalies, where no predecessor product exists, we show results for V3 only. V3 predicts disease density (colonies/75m2) for the GBR (top row) and disease prevalence (percent of host colonies exhibiting signs of disease) for the entire U.S. Pacific (bottom row). The V3 product shows the 75th quantile predicted risk (points) and 50th - 90th quantile predictions (lines). V2 predicts risk levels based on Hot Snap values (units = ℃-weeks, range = 1-15). The validation data shown in these plots were not used in model creation or training.

### Lead time

Both accuracy and precision improved as lead time decreased, but not as drastically and consistently as we expected (Fig. 3), indicating that as the survey date approaches the predictions improve slightly. Positive values for accuracy indicate an over-prediction of disease in the forecast (as shown for white syndromes on the GBR), whilst negative values indicate under-prediction (growth anomalies in both regions). Accuracy improved with shorter lead time for predictions in the GBR for both diseases, while there was almost no improvement with lead time for predictions in the U.S. Pacific. In contrast, precision was largely unaffected by lead time, with marginal improvements for white syndromes in the GBR and growth anomalies in the U.S. Pacific. Given that the variability in SST forecasts decreased with increasing lead time, these results suggest that the prediction uncertainty is largely a function of model uncertainty rather than SST forecast uncertainty.

**Fig. 3.**
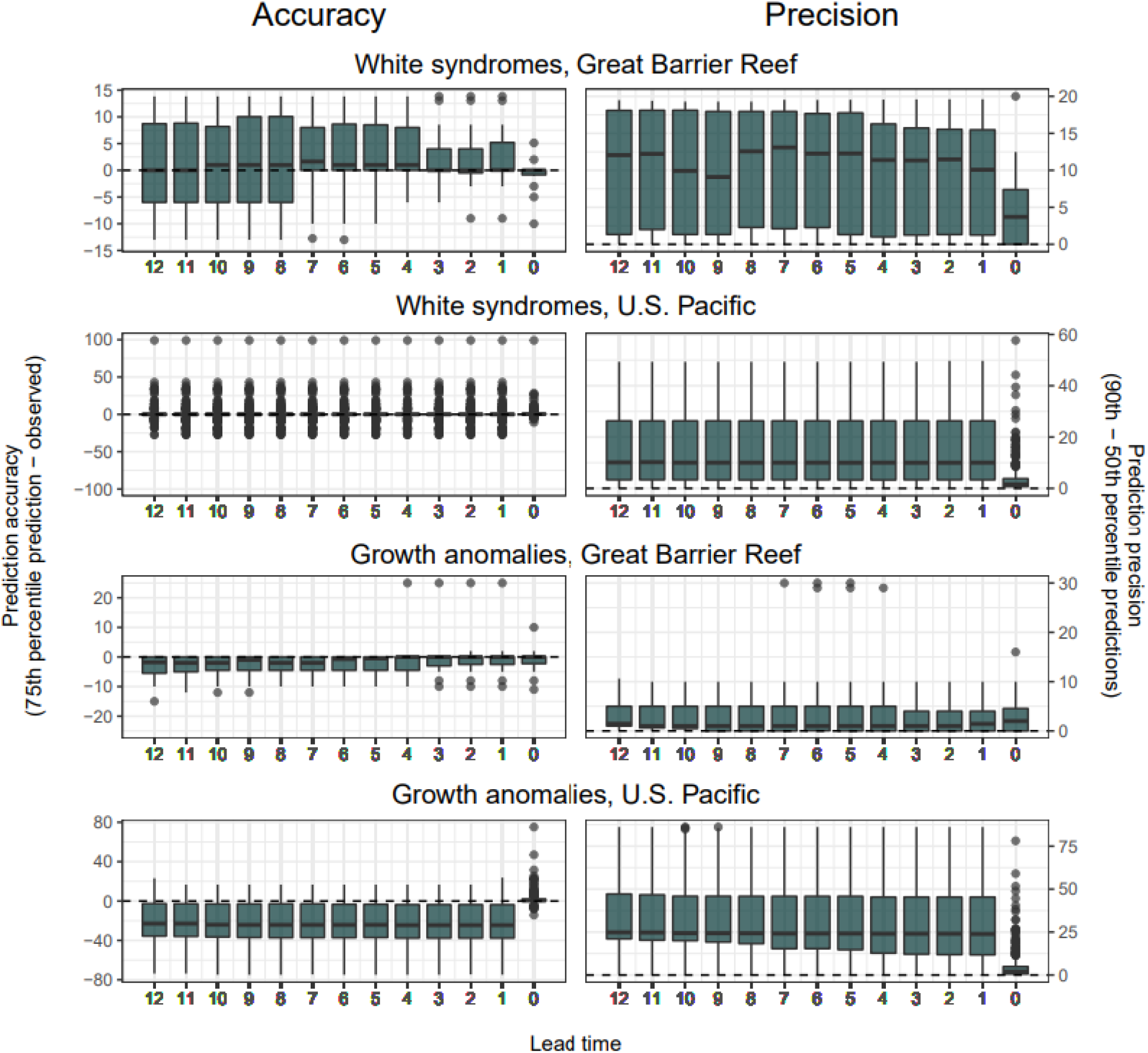
Lead time-dependent predictive accuracy and precision of forecasts. Barplots show predictive accuracy (left column; calculated as difference between 75th quantile prediction and observation) and predictive precision (right column; calculated as difference between 90th and 50th quantile predictions) with different lead times (0-12 weeks prior to observation date). In these plots, perfect accuracy and precision marked by horizontal dashed lines indicate zero difference. Results are shown in eight panels for each of the paired disease types (white syndromes and growth anomalies) and regions (GBR, Australia and U.S. Pacific). Predictions (y-axes) calculated as disease density (colonies/75m2) for the GBR and disease prevalence (percent of host colonies exhibiting signs of disease ranging from 0-100%) for the U.S. Pacific. For example, a median value of 10 for the GBR would indicate that, on average, the model predicts 10 more colonies as diseased than were observed. Similarly, a median value of -20 in the Pacific would indicate that, on average, the model underpredicts disease prevalence by 20%. The validation data shown in these plots were not used in model creation. Month, seasonal turbidity, and SST metrics varied with lead time (in weeks), while all other predictor variables stayed the same (e.g., benthic characteristics of site).

### Coral disease drivers

The most influential disease drivers were primarily time-invariant or seasonally-changing predictor variables (Table 1), which may explain why the V3 product predicts disease with relatively high accuracy for observations from a range of locations and years (Fig. 2), but those predictions do not change substantially with changing lead-times (Fig. 3). The most parsimonious models for each disease-by-region pair varied slightly from each other but broadly reflected relationships found in the literature (Appendix 1: Figs. S1-4). In short, both diseases were primarily influenced by temperature and water quality, coral cover or size, and fish density. White syndromes were strongly influenced by seasonal conditions while growth anomalies were more strongly driven by chronic conditions. A major contribution of this study is the inclusion of multiple metrics of chronic and seasonally changing water quality, which have been shown to influence disease risk in both small-scale correlative and experimental studies (Haapkylä et al. 2011; Pollock et al. 2014; Vega Thurber et al. 2014; Yoshioka et al. 2016), but to-date, have not been possible to include in large scale studies. Thus, this research demonstrates a consistent influence of water quality on disease risk across a broad geographic region and two disease types. Fish density and winter condition were the best predictors of white syndromes in the GBR, followed by variation in seasonal turbidity, summer thermal condition, and coral cover. For white syndromes in the U.S. Pacific, median colony size and chronic and seasonal turbidity metrics (both median and variability for each) were most important. Predictor variables for growth anomalies in both regions were similar to each other, and included 90-day SST mean, fish density, benthic cover metrics, and seasonal and chronic water quality. Within-site water quality variability was more important for growth anomalies in the GBR, whereas average water quality conditions along with coastal development were more important in the U.S. Pacific.

The model selection process revealed that the predictor variables used are better suited for differentiating between lower and higher levels of disease risk rather than presence-absence. We found that the models were able to predict the gradient of observed disease risk best when oversampling surveys in the SMOTE balancing process with relatively high levels of disease risk. For white syndromes, oversampling surveys with >10 diseased colonies/75m^2^ in the GBR and >10% disease prevalence in the U.S. Pacific was optimal; for growth anomalies, oversampling surveys with >15 diseased colonies/75m^2^ in the GBR and >20% disease prevalence in the U.S. Pacific models was optimal (Appendix 1: Fig. S5).

### Decision support tools

The experimental Multi-Factor Coral Disease Risk Forecast, a new tool within NOAA CRW’s decision support system for coral reef management, provides a regional ecological nowcast and forecast of white syndromes and growth anomalies for multiple locations in the Pacific Ocean. Via an online interface on the CRW website (https://coralreefwatch.noaa.gov/product/disease_multifactor/index.php), users can access and explore coral disease forecasts for their region of study, management, and/or interest in the Pacific, to prepare for, monitor, and respond to elevated coral disease risk (Appendix 1: Fig. S6).

### Data explorer

To allow users to explore near real-time, weekly, and seasonal disease predictions more closely, we produced an interactive data explorer tool to complement the NOAA CRW Multi-Factor Coral Disease Risk Forecast. Users can access the data explorer through https://coralreefwatch.noaa.gov/product/disease_multifactor/index.php or at https://coraldisease.com. The data explorer has four components: 1) a disease risk page visualizing nowcasts and forecasts across time and space (Fig. 4); 2) a scenarios page where users can adjust environmental conditions to assess corresponding changes in the nowcast of spatially-explicit disease risk (Appendix S1: Fig. S7); 3) a historical data page that provides information about survey data used to build the models; and 4) an information page with explanatory information and additional resources. Users can explore forecasts and scenarios at multiple spatial scales, ranging from an individual ∼5 km reef pixel to various management zones (containing multiple reef pixels).

**Fig 4.**
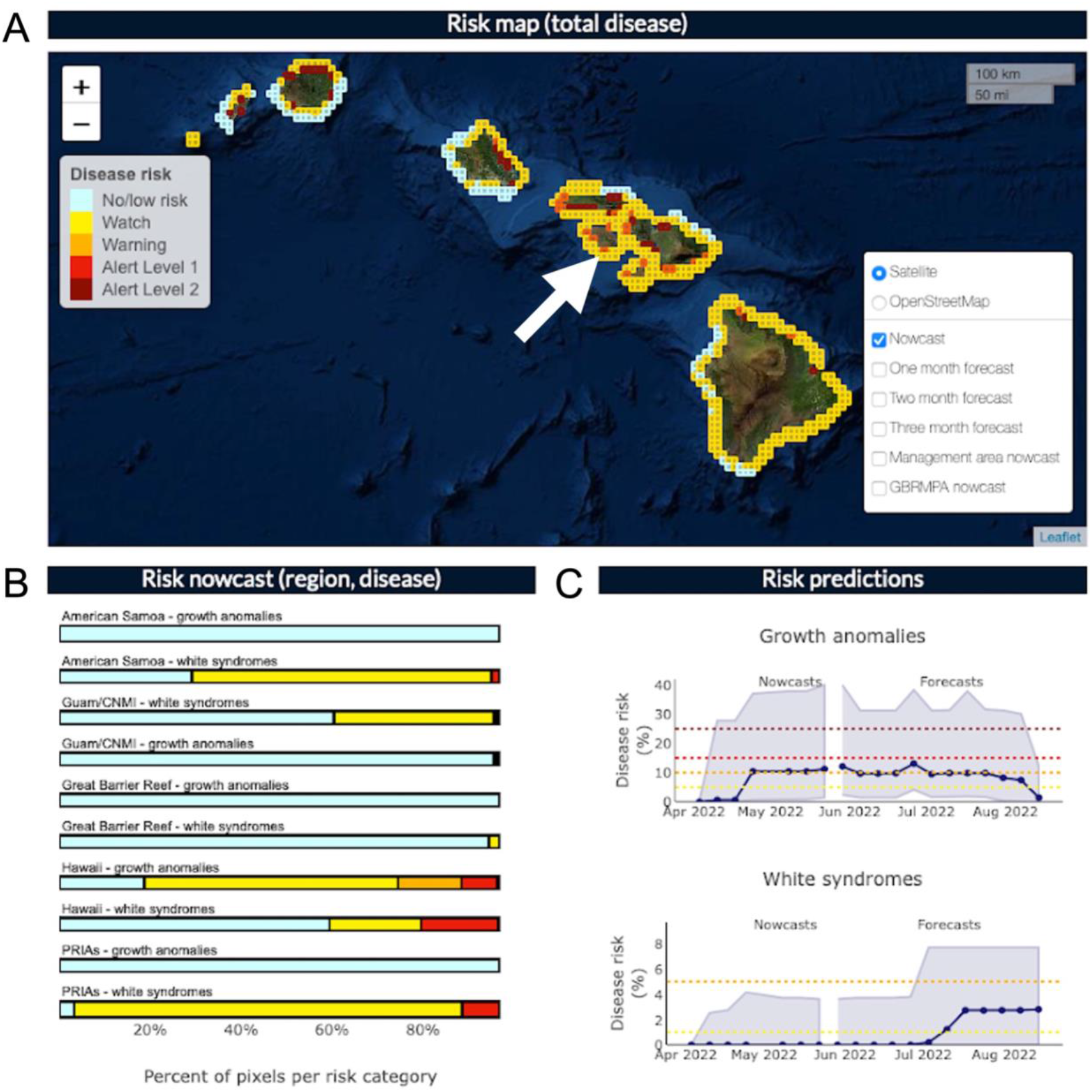
**Data explorer for Multi-factor Coral Disease Risk product**, accessed on 23 May 2022. A) Spatial view of overall color-coded disease risk nowcast for the Main Hawaiian Islands. The thresholds that separate disease risk levels vary by region and disease type (Appendix S1: Table S1). B) Nowcast risk summary for geographic regions and diseases assessed. C) Pixel-specific time-series of nowcasted and forecasted risk on the south coast of Lanai (white arrow in panel A) for growth anomalies and white syndromes, over a 5-month time period.

## Discussion

The Multi-Factor Coral Disease Risk product (V3) offers many improvements over its predecessors, providing a more holistic assessment of disease risk for reefs throughout the Pacific Ocean. In addition to expanding the geographic scope and types of diseases assessed, V3 provides weekly-updated nowcasts and forecasts with up to three months of lead time. The predecessor products fundamentally differed in their forecasting approach; V1 and V2 provide winter pre-conditioning risk outlooks at the end of winter based on wintertime metrics derived from satellite remote sensing data, and then for pixels that are pre-conditioned for risk, refined near real-time predictions are based on satellite monitoring of Hot Snap accumulation throughout the summer months. Thus, within the summer, these prior products produce nowcasts and do not make future predictions; the only prediction component is for the following summer and only at the conclusion of a winter season based on thermal conditions from the entire winter.

Operationally, V3 requires constructing regular predictions of SST-based metrics from climate models rather than relying entirely on near real-time satellite remote sensing (as in V1 and V2). The three-month lead time in V3 aims to provide local stakeholders with more time to organize and execute a response to potential elevated disease risk. The accuracy and precision of disease risk forecasts demonstrate a marginal level of bias in applying the data-based model relationships with predicted values, which may result from variable skill in predicting the inputs (which here are the temperature-based metrics) rather than in the model itself (see further discussion below). Through the online dashboard, users can vary current or predetermined environmental conditions to refine disease risk predictions to better reflect local conditions within the data grid and/or to assess impacts of potential interventions. The most fundamental difference between V1/V2 and V3 is that the new product assesses disease risk based on a suite of ecological conditions in combination with temperature conditions, rather than temperature alone. Some of these new variables such as turbidity were previously unavailable before the incorporation of VIIRS data into these models. Given the relative importance of these new predictor variables (Table 1), we can conclude that although suitable temperature conditions are necessary for elevating risk of white syndromes and growth anomalies, other conditions like colony size and water quality are important driving factors. As a result, the new models that consider a suite of conditions, alongside temperature, have demonstrated better performance in retrospectively predicting disease risk in both the GBR and U.S. Pacific.

While this analysis demonstrates that a suite of conditions are associated with white syndromes and growth anomalies, challenges in forecasting these predictor conditions directly limits capacity for disease prediction. The only predictor variables that are truly forecasted in the Multi-Factor Coral Disease Risk product are the SST-based metrics. For all other variables, we created seasonal climatologies, or rely on time-invariant layers based on long-term aggregated data. For most of the time-invariant variables, such as coral cover, fish densities, and coastal development, we do not expect conditions to change regularly. However, a single event can drastically change biotic conditions on a reef (e.g., a mass bleaching event) and such changes would not be reflected in the forecasts with the predetermined conditions, although they may be assessed (at least to some degree) through adjusting scenarios based on updated information. We anticipate the data may be updatable every 5 to 10 years. We foresee a similar issue for water quality metrics: while we expect that the seasonal climatology and associated variability metric used in these models are fairly robust in the long-term, the current models do not capture acute events caused by intense rainfall and associated runoff, which are known to influence disease (Haapkylä et al., 2011). Although we attempted to measure acute events with ocean color data (procedure described in Geiger et al., 2021), we found that the available data were too sparse to use in the models, with no satellite coverage for ∼80% of the corresponding survey data. More importantly, the ocean color data unavailable during events were not random, but aggregated during cloudy days; in other words, days that are most likely associated with rain events that can increase disease risk. An alternative approach to forecasting water quality conditions could be to create a model based on precipitation forecasts. However, precipitation forecasts are less skillful than temperature forecasts and would require accurate prediction of the timing, intensity, and location of rainfall at fine scales, which must be incorporated into fine-scale hydrologic models with accurate topography and well-predicted initial surface conditions (i.e., soil moisture). Such fine scale hydrologic modeling is generally lacking for most tropical coasts. For this reason, seasonally varying water quality climatologies are the most reliable measurements currently available for coastal coral reefs and applicable for our models. However, we see this process as analogous to early temperature forecasts, which began as almanacs of past conditions (climatologies) and now show high prediction skill through the deployment of increasingly sophisticated statistical and dynamical models.

The extent to which temperature-based metrics are influential in the models determines how well predictions reflect spatial and temporal variability in disease risk. For white syndromes on the GBR for example, both Winter Condition and Hot Snaps are relatively influential variables. As a result, in the retrospective analysis, accuracy and precision varied spatiotemporally – and improved with shorter lead times (consistent with the performance of predicted temperature). In contrast, white syndromes in the U.S. Pacific are less strongly driven by any of the temperature metrics tested in this study, and therefore variability in disease risk is more apparent spatially than temporally. It is worthwhile to note that several white syndromes outbreaks in the U.S. Pacific have occurred in winter (Aeby et al., 2016; Caldwell et al., 2018; Greene et al., 2023; Williams et al., 2011), suggesting that other factors may be more important than temperature in this region and/or that some aspect of temperature not captured by the metrics used in this study is important. For all disease-region pairs, particularly those with less reliance on temperature-based metrics, developing and/or improving climatologies and forecast variables other than temperature would be the most effective way to improve predictability within this forecasting system. A complementary and useful way of leveraging information from V3 is to explore the spatial variability in disease risk to identify locations that are most promising for interventions to improve reef health and target interventions to the most influential variables. For instance, for white syndromes on the GBR, fish density and seasonal turbidity variability were identified as some of the most important predictor variables, indicating that interventions directed at those factors may be most effective for improving reef health. From this perspective, users can explore spatial variability in disease risk and then track any intervention-associated improvements through time without concern over ephemeral conditions that will elapse with weekly updating.

Ecological forecasting presents a variety of ways scientists, managers, and decision-makers can address the rising number of ecological challenges. We provide multiple pathways to explore model predictions and suggest that major improvements going forward will be as dependent on understanding the biological relationships as they are on additional monitoring and surveillance data. The model outputs and associated online Multi-Factor Coral Disease Risk product and data explorer were co-developed with many relevant management agencies and scientists. Through multiple focus groups with stakeholders in Australia, American Samoa, Hawaii, and Guam, planning meetings, and workshop demonstrations at several scientific conferences over the course of six years, we created an online decision support tool that provides regional overviews aligned with other NOAA CRW tools with which our intended audience is already familiar. The method of delivering regional overviews is also preferred by users with slow or intermittent internet connection, as is common in some Pacific islands. The data explorer complements this tool in several ways. First, it provides predictions aggregated to relevant management zones and allows users to explore forecasts at these various spatial scales through time. This addresses two key concerns of our users as they need to distill information at scales relevant to their respective agencies or work mandates, and to understand trends through time in those specific locations. We addressed a suite of other concerns through the use of scenarios. Broadly, users who interacted with the tools as they were being developed and tested found it difficult to translate mean conditions at the finest spatial scale (∼5 km) available to an individual reef of interest, especially when they knew conditions at that one location were different from the surrounding region. Thus, we made it possible for users to change individual input conditions in the scenarios page of the interactive tool to see how predicted disease risk may correspondingly change in a specified area of interest. The same scenarios tool can alternatively be used as an exercise to assess the predicted impacts of an intervention that would affect the relevant input conditions (e.g., an intervention to reduce resuspended sediments via turbidity) to determine how that might affect disease risk.

Going forward, the forecasting models could be substantially improved by replacing phenomenological relationships with biological ones and potentially by calibrating the models differently. Ideally, biological relationships could replace phenomenological ones by using a combination of lab and natural experiments. This approach would ultimately help reduce uncertainty, particularly for undersampled conditions. In terms of calibration, we made several decisions that increased the likelihood of false positives (i.e., predicting higher disease levels than would be observed). Specifically, using SMOTE to compensate for scarce data on the conditions associated with elevated disease risk resulted in an overrepresentation of those conditions in the model data. Further, we used the 75th percentile when communicating the model results in an effort to guard against missing a major disease event. The impact of these decisions plays out as expected with a large number of false positives in the validation exercise (Fig. 2). While we made these choices based on stakeholder input, it might be preferable in future work to calibrate the models in a way that systematically assesses a broader suite of assumptions and allows for optimization of those decisions. For instance, future efforts might include performing a formal parameter sweep across a broader range of SMOTE data frequencies and prediction quantiles. Alternatively, if enough information is known about the disease system, one could use informative priors in a Bayesian analysis or consider adding a base rate correction.

The overall modeling approach we used to create V3 could be replicated to predict disease risk for other reef regions and diseases, with appropriate consideration given to the transferability of input variables to these model systems. To expand this framework, a model would need to be developed tailored to the new location and/or disease. This would require the collation of coral health survey data and concurrent environmental conditions for model development, and collating gridded environmental covariates, including climatologies and SST forecasts, for the appropriate reef grid for forecasting. Diseases most suited for a forecasting framework like the one described in this study are those impacting widely distributed hosts, where the burden shifts seasonally between endemic and epizootic states. For example, Stony Coral Tissue Loss Disease has caused widespread mortality in multiple species in the Caribbean and would be an ideal candidate disease for expanding the current framework if it were introduced to the Pacific basin or through the expansion of this tool to the western Atlantic.

Many of the issues that make it challenging to forecast coral disease risk are issues that encumber ecological forecasts more broadly. In many ecological systems, the greatest obstacle is data limitation. For example, while we had extensive coral disease survey data that spanned a large geographic range in the Pacific Ocean and a broad time horizon, very few of the data points contained useful information about disease density or prevalence, as most surveys exhibited low or disease-free conditions. This problem is likely to arise in other attempts to forecast low occurrence events such as infestations, invasive species, tipping points, and extreme events. While the historical low occurrence of disease is good ecologically, these data limitations inhibit both our ability to develop initial ecological forecasts and to create a workflow with continual validation and updates, which has been key to improving forecasts in other systems such as weather, storm, and fire forecasting (Dietze et al., 2018). A complementary issue is the reliance on forecasted data as inputs to an ecological forecasting model, which may have their own set of uncertainties and challenges. An important question then arises from these shared obstacles across systems: is there something inherently different and currently unknown about developing forecasts in systems where data cannot be regularly updated and validated? Thus, this ecological forecast and many others will benefit from community-wide progress in the field of ecological forecasting.

## Conclusions

Herein, we present the next-generation NOAA CRW coral disease forecasting product and an associated data explorer tool. It provides many advantages over its predecessors, including near-term forecasts of coral disease risk in many major reefs in the Pacific Ocean. The Multi-Factor Coral Disease Risk product predicts disease risk for white syndromes and growth anomalies with greater precision and accuracy than previous products based on temperature alone, and provides information for more diseases and regions. Co-developing the user interface with the intended user base of scientists and managers resulted in a user-friendly online data explorer tool that includes assessment of disease risk at different scales, quantification of uncertainty in predictions, and the ability to adjust input conditions to assess effects on disease outcomes. While this iteration is a major improvement to the NOAA CRW coral disease forecasting products, largely thanks to numerous advances in the ecological forecasting community and data availability, there are still numerous limitations for forecasting coral disease risk. As data availability, forecasting capabilities, and our biological understanding of the system improves, so can future versions of this product.

## Acknowledgments

Sincere thanks to Maury Estes of the NASA Ecological Forecasting Program for guidance and support throughout the project. Thanks also to the numerous people who provided feedback during the development of the tools through formal workshops, user-engagement meetings, and/or personal input as well as constructive feedback from two anonymous Reviewers. The scientific results and conclusions, as well as any views or opinions expressed herein, are those of the authors and do not necessarily reflect the views of NOAA or the Department of Commerce.

