## Appendix_S1 for "Multi-Factor Coral Disease Risk Forecasting for Early Warning and Management"

*Ecological Applications*

**Table S1. Warning levels.** There are five warning levels of disease risk displayed on the maps and graphs in the decision support tool, reflecting predicted disease severity: Low Risk, Watch, Warning, Alert Level 1, and Alert Level 2. The thresholds separating warning levels were determined based on historical disease levels and expert elicitation, and vary by disease type and region.

| Warning level | Growth anomalies Australia | Growth anomalies U.S. Pacific | White syndromes Australia | White syndromes U.S. Pacific |
|---|---:|---:|---:|---:|
| **Low risk** | 0-5 colonies | 0-5% | 0-1 colonies | 0-1% |
| **Watch** | 6-15 colonies | 6-10% | 2-5 colonies | 2-5% |
| **Warning** | 16-25 colonies | 11-15% | 6-10 colonies | 6-10% |
| **Alert level 1** | 26-50 colonies | 16-25% | 11-20 colonies | 11-15% |
| **Alert level 2** | >50 colonies | >25% | >20 colonies | >15% |

**Table S2.** Hypothesized predictor variables and their data sources. ✅ indicates the variable was tested in the disease-by-region model. Acronyms: GBR = Great Barrier Reef; WS = white syndromes; GA = growth anomalies; AIMS LTMP = Australian Institute of Marine Science Long Term Monitoring Program; NOAA NCRMP = National Oceanic and Atmospheric Administration National Coral Reef Monitoring Program; NASA VIIRS = National Aeronautics and Space Agency Visible Infrared Imaging Radiometer Suite (VIIRS) instrument aboard the Suomi-NPP satellite; CRW = NOAA Coral Reef Watch.

| | | White syndromes | | Growth anomalies | |
|---|---|---|---|---|---|
| Variable | Definition (source) | GBR | U.S. Pacific | GBR | U.S. Pacific |
| *Time-invariant predictors* | | | | | |
| Coral cover | Total live coral cover, specific to morphology or family if indicated below [range = 0-100%] (surveys)<br>● Plating and table [WS, GBR]<br>● Acroporidae [WS, U.S. Pacific]<br>● Poritidae [GA, U.S. Pacific] | ✅ | ✅ | ✅ | ✅ |
| Median colony size | Site-specific median colony size of corals in the family Acroporidae [WS] or Poritidae [GA] (surveys) | | ✅ | | ✅ |
| Colony size variability | Site-specific coefficient of variation of colony size of corals in the family Acroporidae [WS] or Poritidae [GA] (surveys) | | ✅ | | ✅ |
| Herbivorous fish density | GBR: Herbivorous fish count (per ~2 km) based on manta tow surveys between 1986-2005. (AIMS LTMP)<br><br>U.S. Pacific: Herbivorous fish density | ✅ | ✅ | ✅ | ✅ |

| | | | | | |
|---|---|---|---|---|---|
| | (per $m^2$) by reef sectors averaged from surveys collected between 2007-2012 (NOAA NCRMP) | | | | |
| Parrotfish density | Parrotfish density (per $m^2$) by reef sectors averaged from surveys collected between 2010-2019 (NOAA NCRMP) | | ✅ | | |
| Butterflyfish density | Butterflyfish density (per $m^2$) by reef sectors averaged from surveys collected between 2010-2019 (NOAA NCRMP) | | ✅ | | |
| Long term Kd(490) median | Site-specific long-term median satellite Kd(490) ($m^{-1}$) from 2012-2020 within a 5-pixel buffer (NASA VIIRS) | ✅ | ✅ | ✅ | ✅ |
| Long term Kd(490) variability | Site-specific variability of weekly satellite Kd(490) ($m^{-1}$) from 2012-2020 within a 5-pixel buffer (NASA VIIRS) | ✅ | ✅ | ✅ | ✅ |
| Coastal development | Artificial light intensity (high gain) (0-255) at 3 km resolution (NASA Black Marble 2016 dataset: https://blackmarble.gsfc.nasa.gov/) | | | ✅ | ✅ |
| *Seasonally-changing predictors* | | | | | |
| Three-week Kd(490) median | Site-specific median of three week moving window of satellite Kd(490) ($m^{-1}$) from 2012-2020 based on a 5-pixel buffer [i.e., values from week before, during, and after survey] (NASA VIIRS) | ✅ | ✅ | ✅ | ✅ |
| Three-week Kd(490) variability | Site-specific variability for three week moving window of satellite Kd(490) ($m^{-1}$) from 2012-2020 based on a 5-pixel buffer [i.e., values from week | ✅ | ✅ | ✅ | ✅ |

| | | | | | |
|---|---|---|---|---|---|
| | before, during, and after survey] (NASA VIIRS) | | | | |
| Month | Month of year (surveys) | ✓ | ✓ | ✓ | ✓ |
| *Regularly-changing predictors* | | | | | |
| 90-day SST mean | Mean sea surface temperature for 90 days prior to survey date (°C) (CRW CoralTemp) | | | ✓ | ✓ |
| Hot Snap | Accumulated heat stress combining magnitude and duration of SST above a locally-specific summertime threshold at 5 km resolution (°C) (CRW CoralTemp) | ✓ | ✓ | | |
| Winter Condition | Accumulated thermal conditions combining magnitude and duration of positive and negative SST anomalies about the locally-specific winter average, including all dates within the climatological three-month winter period and any temperatures cooler than one standard deviation above the winter average (°C) (CRW CoralTemp) | ✓ | ✓ | | |

**Table S3.** Data sources and methodology for coral disease surveys.

| Data source | Sampling methodology | Site selection | Reference(s) |
|---|---|---|---|
| NOAA NCRMP | Belt transect | Stratified random | https://doi.org/10.7289/v53n21q5 https://doi.org/10.7289/v579431k https://doi.org/10.7289/v5c24trh https://doi.org/10.7289/v5zw1j8b |
| University of Guam | Belt transect | Fixed sites | Author contributed (Raymundo) |
| Hawaii Coral Disease database | Belt transect, Line Point Intercept | Predominantly stratified random | Caldwell, Burns et al. 2016 |
| Great Barrier Reef Marine Park Authority | Reef health impact surveys | Mixture of random, responsive (to observed impact), and fixed (e.g., tourism operator platforms) | Beeden et al. 2014 |

**Table S4.** Most common genera and species in each region. Proportion shown in parentheses. CNMI = Commonwealth of the Northern Mariana Islands; MHI = Main Hawaiian Islands; NWHI = Northwestern Hawaiian Islands; PRIA = Pacific Remote Island Area (also called U.S. Pacific Remote Islands Marine National Monument).

| **Region** | **Family** | **Most common genera** | **Most common species** |
|---|---|---|---|
| Guam | Acroporidae | Acropora (95%) | *Acropora pulchra* (63%)<br>*Acropora azurea* (15%)<br>*Acropora muricata* (8%) |
| Guam | Poritidae | Porites (>99%) | *Porites cylindrica* (64%)<br>*Porites rus* (21%) |

| | | | |
|---|---|---|---|
| CNMI | Acroporidae | Astreopora (54%) | Not specified (65%)<br>*Astreopora myriophthalma* (29%) |
| | | Montipora (32%) | Not specified (89%) |
| CNMI | Poritidae | Porites (94%) | Not specified (65%)<br>*Porites rus* (15%)<br>*Porites lobata* (12%) |
| MHI | Acroporidae | Montipora (100%) | Not specified (54%)<br>*Montipora capitata* (46%) |
| MHI | Poritidae | Porites (100%) | *Porites lobata* (78%)<br>*Porites compressa* (13%) |
| NWHI | Acroporidae | Montipora (80%) | *Montipora capitata* (54%)<br>*Montipora flabellata* (23%)<br>*Montipora patula* (19%) |
| | | Acropora (20%) | *Acropora cytherea* (79%)<br>Unspecified (17%) |
| NWHI | Poritidae | Porites (100%) | *Porites lobata* (58%)<br>*Porites compressa* (16%)<br>*Porites lichen* (10%) |
| PRIA | Acroporidae | Montipora (81%) | Unspecified (62%)<br>*Montipora foveolata* (19%)<br>*Montipora aequituberculata* (9%) |
| | | Acropora (17%) | Unspecified (66%)<br>*Acropora nobilis* (17%)<br>*Acropora cytherea* (8%) |
| PRIA | Poritidae | Porites (100%) | *Porites vaughani* (35%)<br>Unspecified (25%)<br>*Porites lobata* (21%) |
| American Samoa | Acroporidae | Montipora (76%) | Unspecified (85%)<br>*Montipora foveolata* (8%) |

| | | | |
|---|---|---|---|
| | | Acropora (12%) | Unspecified (92%) |
| | | Astreopora (9%) | Unspecified (60%)<br>*Astreopora myriophthalma* (40%) |
| American Samoa | Poritidae | Porites (95%) | *Porites rus* (34%)<br>Unspecified (34%)<br>*Porites lichen* (12%) |

**Table S5.** Data from disease surveys were highly unbalanced, with the majority of surveys reporting no disease. Number of surveys shown in parentheses.

| | White syndromes | | Growth anomalies | |
|---|---|---|---|---|
| | GBR | U.S. Pacific | GBR | U.S. Pacific |
| Disease-free surveys (N) | 97% (35,352) | 89% (1,558) | 97% (34,919) | 77% (1,415) |
| Disease-present surveys (N) | 3% (1,203) | 11% (169) | 3% (1,160) | 33% (324) |

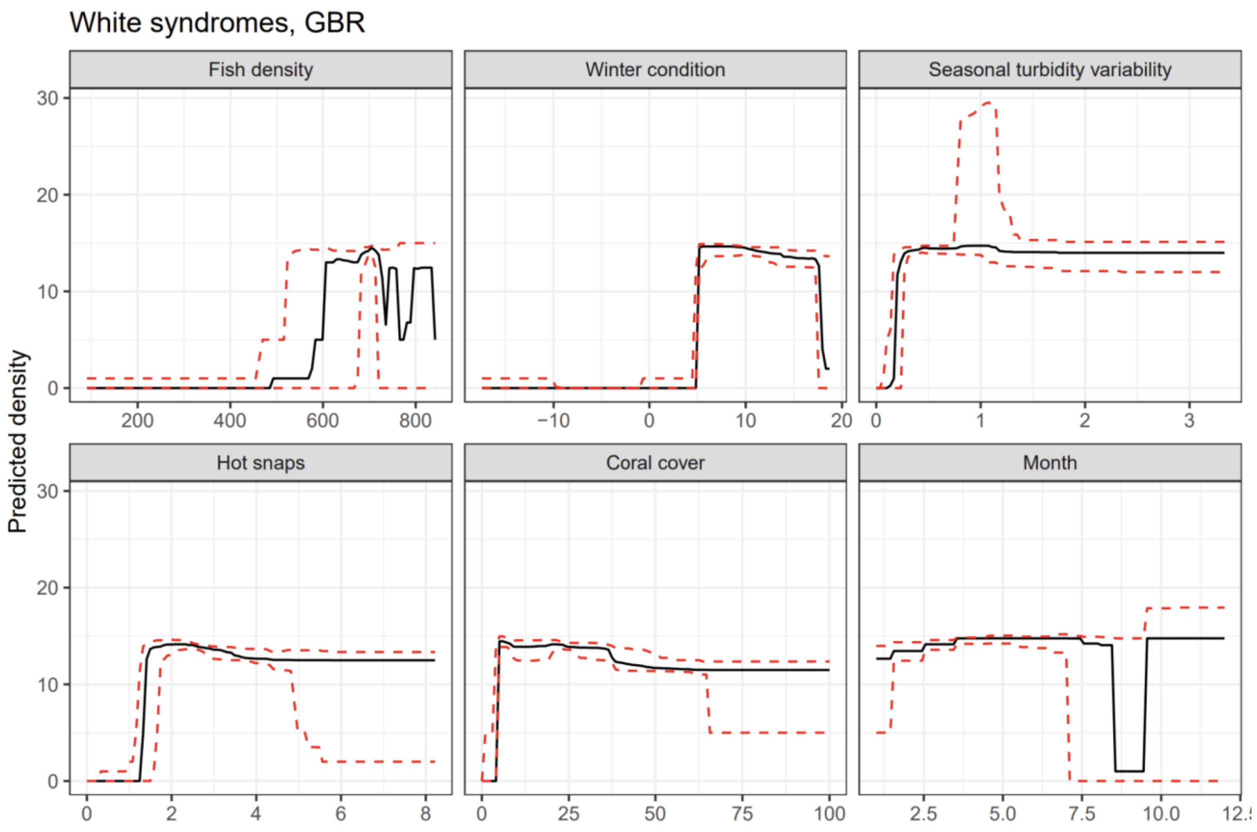


**Fig S1. Partial dependence plots visualizing relationships between predictor variables and white syndromes disease risk for the east coast of Australia.** Individual plots show relationships between predicted density of disease colonies/75m$^2$ (y-axes) and the predictor variable labeled at the top of each subplot (x-axes); the axis units vary by predictor variable. The black lines show the 75th quantile, red dashed lines show the 50th and 90th quantiles.

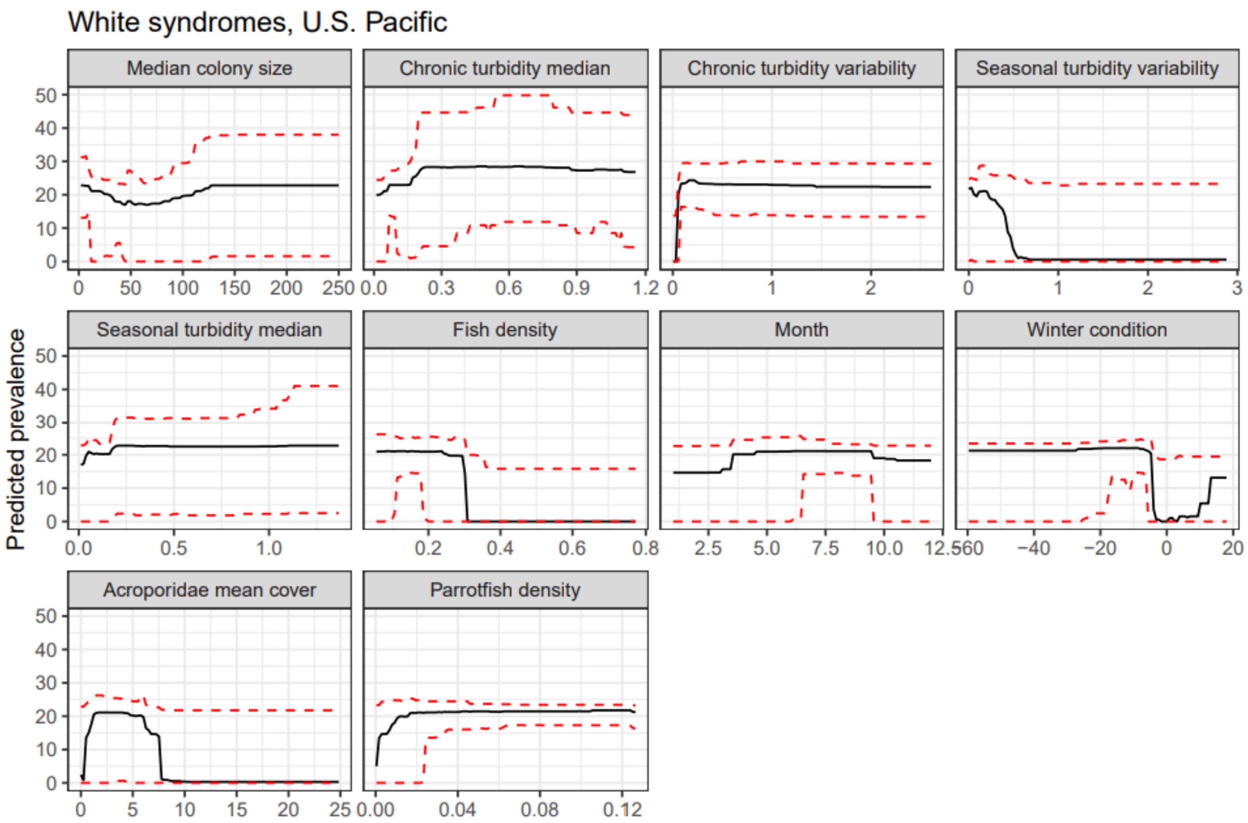


**Fig S2. Partial dependence plots visualizing relationships between predictor variables and white syndromes disease risk for the U.S. Pacific.** Individual plots show relationships between predicted prevalence of disease colonies (y-axes) and the predictor variable labeled at the top of each subplot (x-axes); the axis units vary by predictor variable. The black lines show the 75th quantile, red dashed lines show the 50th and 90th quantiles.

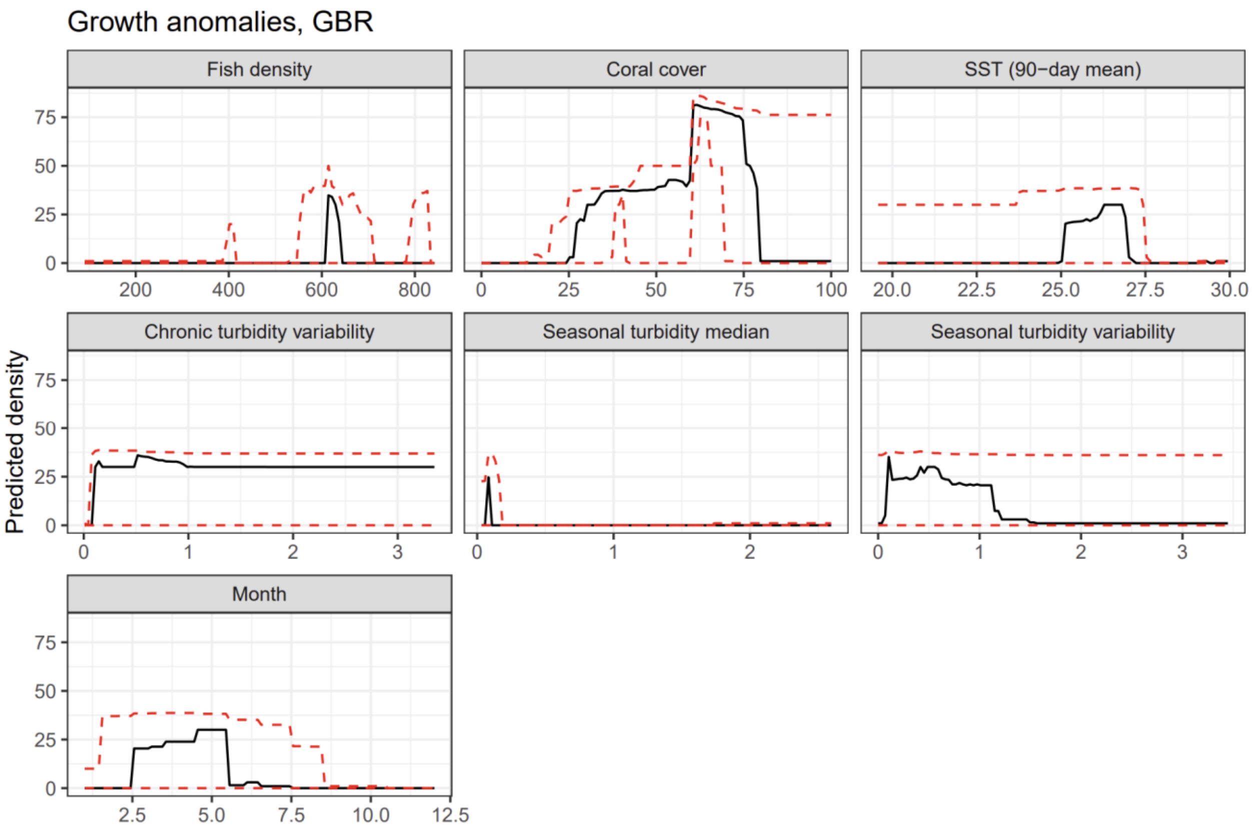


**Fig S3. Partial dependence plots visualizing relationships between predictor variables and growth anomalies disease risk for the east coast of Australia.** Individual plots show relationships between predicted density of disease colonies/75m$^2$ (y-axes) and the predictor variable labeled at the top of each subplot (x-axes); the axis units vary by predictor variable. The black lines show the 75th quantile, red dashed lines show the 50th and 90th quantiles.

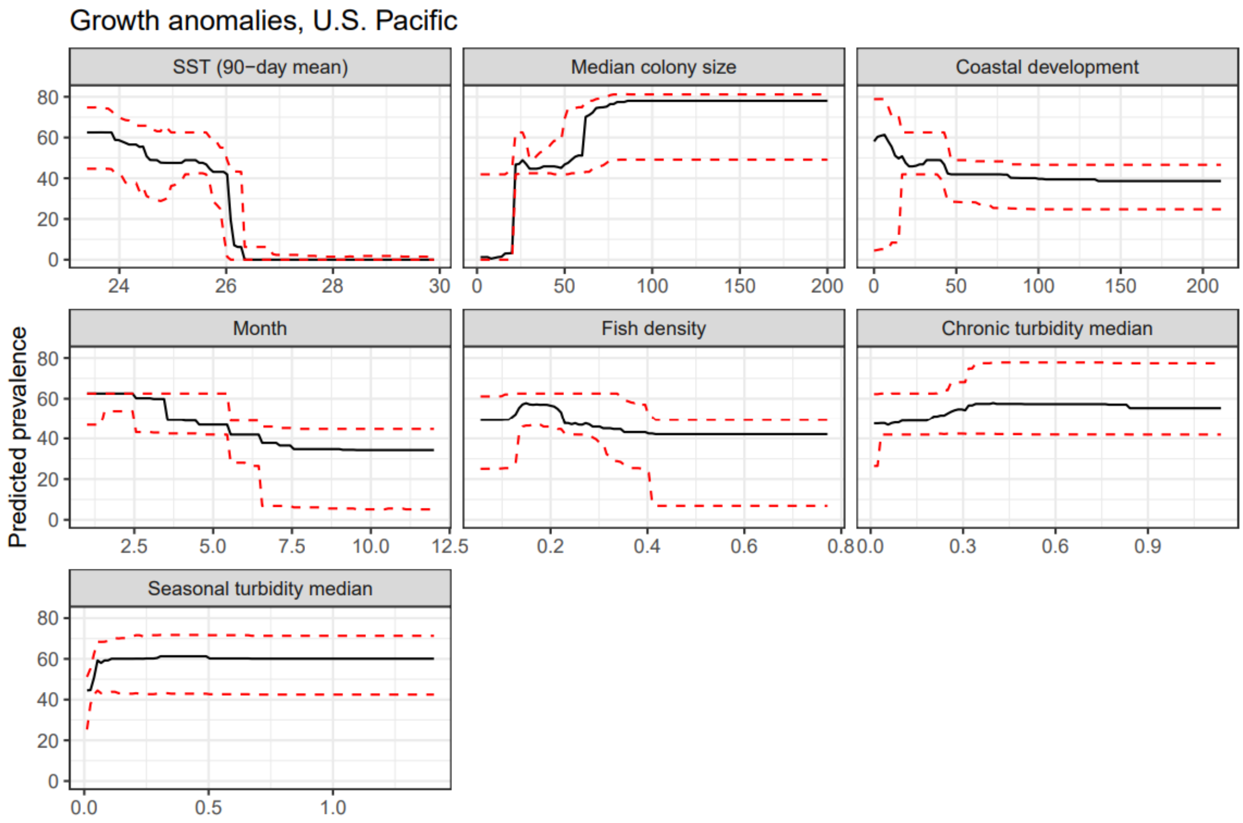


**Fig S4. Partial dependence plots visualizing relationships between predictor variables and growth anomalies disease risk for the U.S. Pacific.** Individual plots show relationships between predicted prevalence of disease colonies (y-axes) and the predictor variable labeled at the top of each subplot (x-axes); the axis units vary by predictor variable. The black lines show the 75th quantile, red dashed lines show the 50th and 90th quantiles.

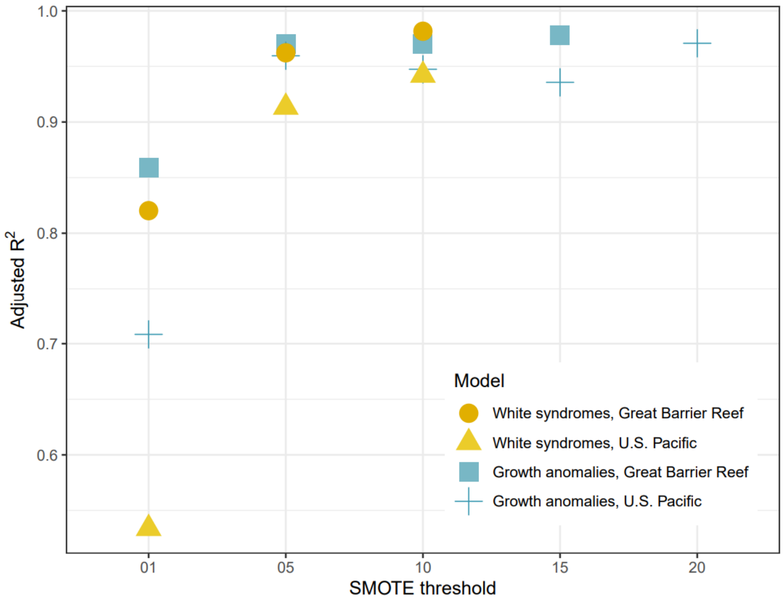


**Fig S5.** Adjusted $R^2$ value for the most parsimonious model per SMOTE threshold across disease-by-region pairs based on withheld data. We selected the model with the overall highest adjusted $R^2$ value from this analysis to use operationally in the Multi-Factor Coral Disease Risk product.

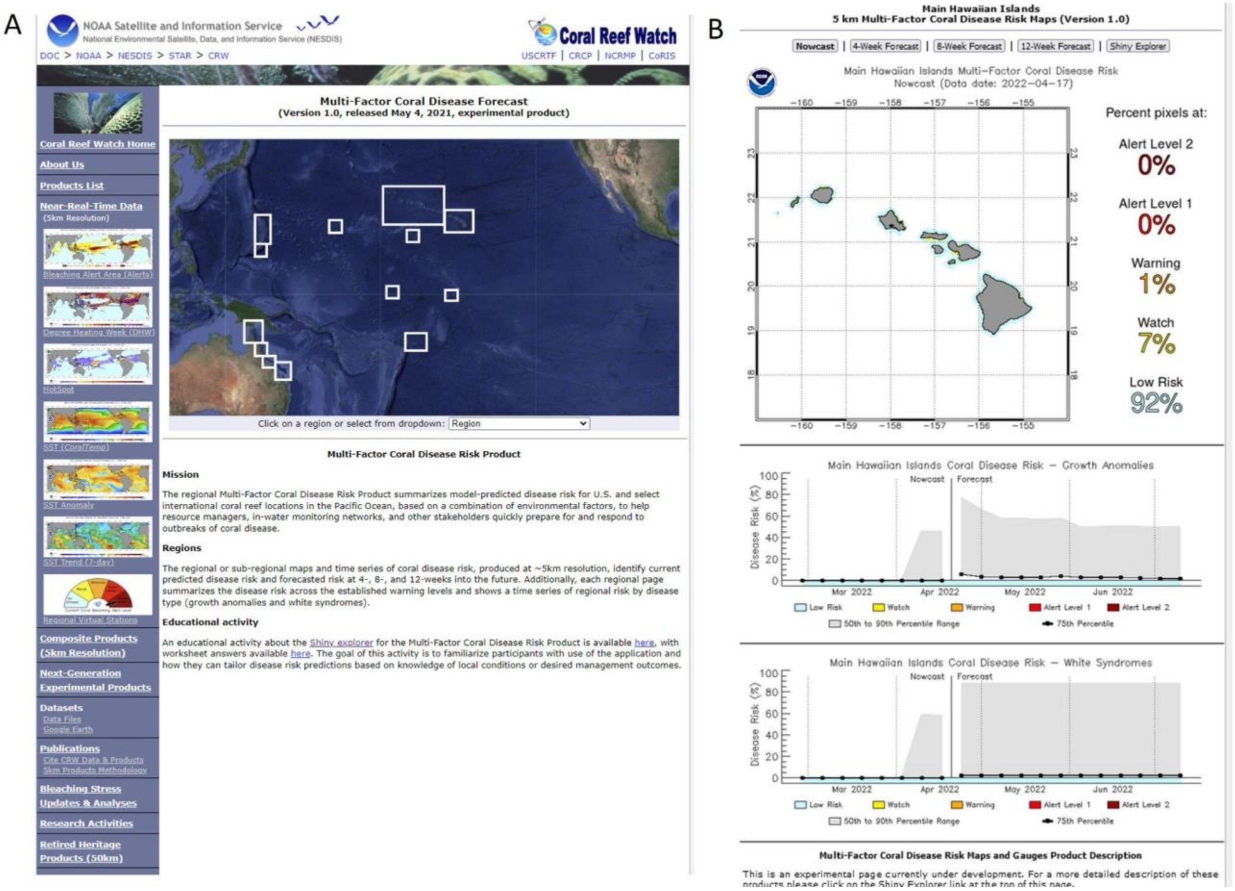


**Fig. S6.** A) Example screenshot of the Multi-Factor Coral Disease Risk product interface on the NOAA Coral Reef Watch website. B) Each region shows a map and numeric summary of risk categories across the region, with disease-specific time series. This example is for the Main Hawaiian Islands. Users can toggle among 4-, 8-, and 12-week forecasts and connect to the data explorer.

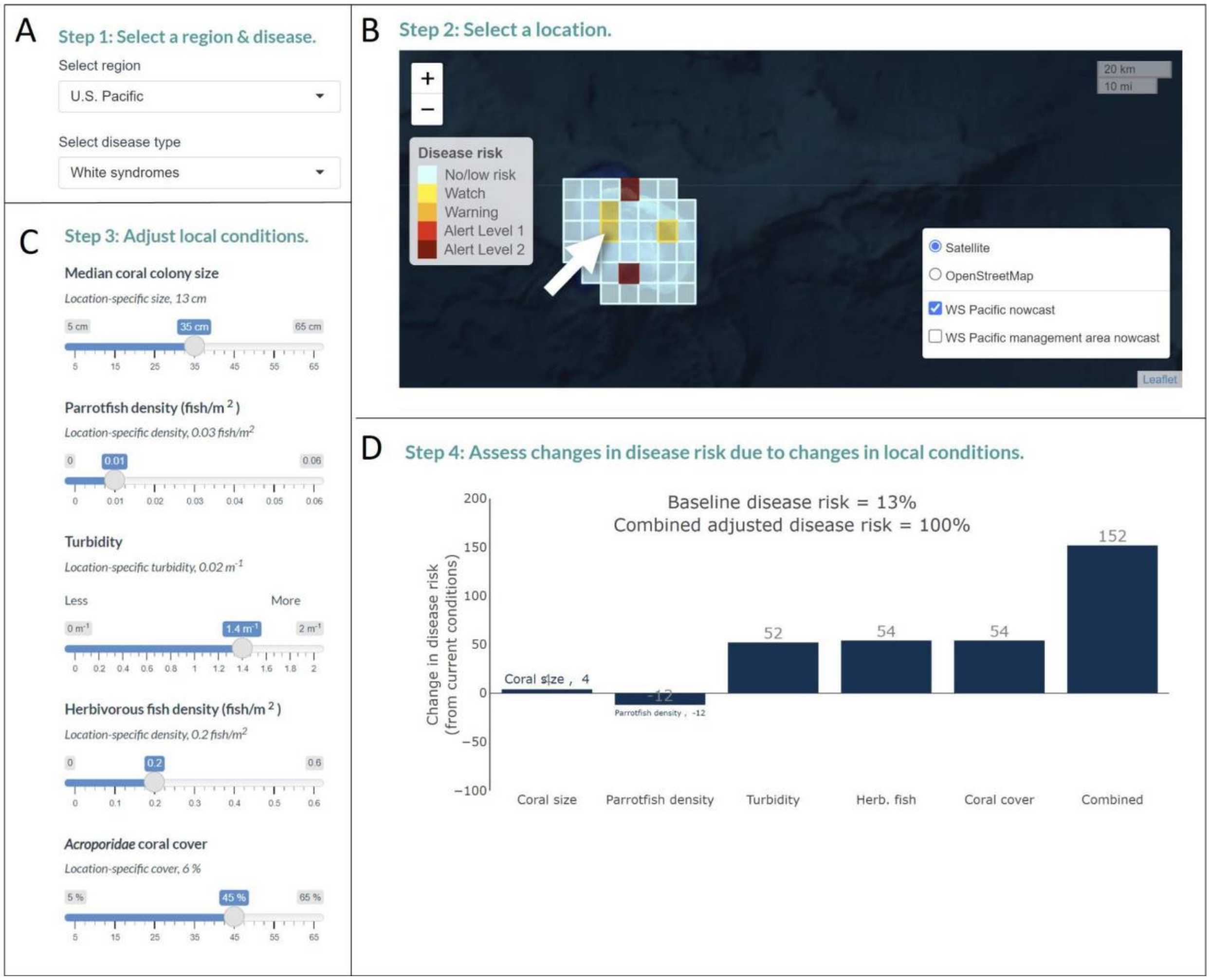


**Fig. S7**. **Example data explorer for Multi-factor Coral Disease Risk product ‘Investigating scenarios’ tab.** A) User selects the region and disease of interest. B) User selects a single pixel or management area to explore. C) Sliders are initially set at the current reef-pixel-specific values for select predictor variables used for producing default weekly-updated disease risk predictions. Users can adjust sliders to better reflect local conditions or assess potential intervention scenarios. D) Plot indicates current (baseline) predicted disease risk as well as adjusted disease risk based on conditions selected (i.e., slider values). Bars indicate change in baseline disease risk associated with each variable.
